# Architecture-Dependent Transition from Quasi-2D to 3D Cell–Scaffold Interactions in Ultrafine Electroprinted Cellulose Scaffolds

**DOI:** 10.64898/2026.09.08.750081

**Authors:** Mahsa Jamadi Khiabani

## Abstract

Precise control over scaffold microarchitecture is critical for engineering cell-instructive biomaterials and for understanding how structural cues regulate cell–material interactions. In this study, ultrafine cellulose-based scaffolds fabricated by near-collector electroprinting (NCE) were used as a model platform to investigate how scaffold architecture influences the transition between quasi-2D and 3D cell–scaffold interactions. Cellulose acetate scaffolds with micrometer-scale fiber diameters, tunable fiber spacing, and defined multilayer geometries were systematically evaluated to determine the effects of scaffold dimensionality and pore architecture on human mesenchymal stem cell behavior. Low-layer scaffolds functioned primarily as quasi-2D topographical patterns, where cells adhered predominantly to the underlying substrate while aligning along the printed fibers in a spacing-dependent manner. Increasing scaffold height generated a 3D microenvironment that promoted direct cell–scaffold interaction and enabled inter-fiber bridging. Quantitative analysis of nuclear orientation and actin organization revealed that scaffold dimensionality and fiber spacing jointly govern the transition between topographical guidance and bridging-mediated cellular organization. To further demonstrate material versatility, lignosulfonate was incorporated into the printing ink to fabricate composite scaffolds while retaining print fidelity and cytocompatibility. Together, these findings demonstrate how scaffold architecture can be used to govern cell–material interactions across multiple dimensional regimes.

## 1. Introduction

The development of advanced tissue scaffolds for bioengineering applications is a rapidly growing field, driven by the need for materials and fabrication methods that offer precise structural control and tailored biological performance^1^. A central aim in tissue engineering and related biomedical applications is to recreate the complexity of native tissue by enhancing both biological and mechanical properties^1–3^. Key scaffold features—such as porosity, pore architecture, and fiber alignment—play a critical role in guiding cellular behavior, influencing cell adhesion, proliferation, and differentiation^4–7^. In particular, surface patterning and microscale topography are key design parameters for regulating cellular functions in engineered scaffolds^8^. This interdisciplinary challenge has fueled innovations at the intersection of materials science, engineering, and biology, leading to the emergence of diverse scaffold fabrication approaches^9^. Among these, electrospinning and 3D printing have gained particular attention for their ability to produce scaffolds with micro- and nano-scale features^10–13^.

Electrospinning is widely adopted for scaffold fabrication due to its ability to produce ultrafine fibers that closely mimic the extracellular matrix^14, 15^. In this technique, a polymer solution is dispensed through a nozzle and exposed to a high-voltage electrostatic field, forming continuous fibers that are collected on a grounded surface^11, 12, 14^. Electrospinning parameters—such as polymer molecular weight, applied voltage, and flow rate—significantly influence fiber properties^11, 16^. However, conventional electrospinning typically yields randomly deposited, sheet-like structures with limited control over fiber placement and minimal capability for vertical (z-axis) structuring. As a result, it is not well suited for the fabrication of architected 3D scaffolds with defined geometry^17^.

To overcome these limitations, additive manufacturing approaches have been developed to enable spatially controlled scaffold fabrication^10, 13, 17^. Advanced 3D printing techniques allow layer-by-layer construction of structures with defined geometry and tunable porosity, supporting mechanical integrity, mass transport, and cell growth^10, 13, 17^. Among these, two-photon polymerization offers sub-micron resolution but is restricted by the choice of materials^18^. Extrusion-based techniques such as fused deposition modeling and direct ink writing (DIW) enable fabrication of scaffolds with pore sizes typically ranging from 100 to 600 µm^19, 20^; however, achieving resolutions below 100 µm remains challenging and strand merging occurs when attempting to print at smaller distances^21, 22^. Although DIW can reach resolutions of ∼30 µm in hydrogel systems ^23^, maintaining such fine features in multilayered structures is difficult, and the use of small nozzles increases clogging risk while restricting material selection^24, 25^. Melt electrowriting (MEW) allows for more precise microstructure fabrication^26^, producing scaffolds with pore sizes as small as 40 µm^27^, extending up to several hundred microns^28, 29^. However, reproducibility becomes a major challenge below 200 µm, as smaller pores are difficult to achieve with high precision and consistency^30, 31^. Additionally, MEW requires the material to be melted, which limits the choice of biomaterials and may compromise the biological functionality of sensitive components^7^.

Electroprinting can be described as bridging the gap between electrospinning and DIW. Like electrospinning, it utilizes electrohydrodynamic forces to draw fibers, but it operates with a much shorter nozzle-to-collector distance, typically in the range of 500 µm to 3 mm^32^, which allows for more controlled fiber placement^33^. This technique enables the fabrication of multilayered scaffolds with well-defined architectures^33, 34^. Despite its advantages, conventional electroprinting faces challenges in achieving fine fiber diameters and maintaining structural fidelity, particularly for well-defined multilayered 3D architectures^35^. This limitation reduces its suitability for applications requiring high-resolution features. To address this limitation, near-collector electroprinting (NCE) has been introduced, in which the nozzle-to-collector distance is reduced to the order of tens of micrometers, enabling enhanced control over fiber placement and the fabrication of scaffolds with pore sizes well below 100 µm, thereby improving structural resolution^36, 37^. In this setup, the electric field is applied over a centimeter-scale distance, while an intermediate insulating capillary assists in stabilizing the ink flow and controlling fiber placement^37^. A key advantage of NCE over DIW is the ability to use larger nozzle openings, thereby reducing clogging risks while maintaining high-resolution fiber deposition. While related techniques such as near-field electrospinning (NFES) have demonstrated controlled deposition of fibers with ∼10 µm spacing, their application has largely been limited to planar architectures^38^. Despite these advances, the application of NCE for biomedical use and cell–material interaction studies remain underexplored.

In this study, ultrafine scaffolds fabricated by NCE were used to investigate how scaffold architecture regulates human mesenchymal stem cell (hMSC) behavior. Cellulose acetate (CA), a cellulose derivative with favorable biocompatibility, mechanical stability, and excellent processability, was selected as the base material for scaffold fabrication^39^. By systematically varying fiber spacing and printed layer number, the influence of scaffold architecture on the viability, alignment, spreading, and cytoskeletal organization of hMSCs was investigated. Particular emphasis was placed on identifying how scaffold dimensionality modulates the transition between quasi-2D topographical guidance and 3D bridging-mediated cell–scaffold interactions. Furthermore, to demonstrate the compositional versatility of the NCE platform, lignosulfonate (LS) was incorporated into CA inks to fabricate lignin-containing composite scaffolds and assess their printability and cytocompatibility. Together, this study provides insight into how scaffold dimensionality and pore architecture can be used as design variables to regulate cell–material interactions in ultrafine fibrous constructs.

## 2. Materials and Methods

### 2.1. Materials

CA (Mw 50 kDa), LS, acetone, dimethyl sulfoxide (DMSO), paraformaldehyde, Triton X-100, bovine serum albumin (BSA), and phosphate-buffered saline (PBS) were purchased from Sigma-Aldrich and used as received. Dulbecco’s Modified Eagle Medium (DMEM), fetal bovine serum (FBS), and penicillin–streptomycin were obtained from Gibco (USA). LIVE/DEAD Viability/Cytotoxicity Kit and Alexa Fluor 594 Phalloidin were obtained from Invitrogen (Thermo Fisher Scientific, Sweden). 4′,6-Diamidino-2-phenylindole (DAPI) was purchased from Merck (Sweden).

### 2.2. NCE Setup

Scaffolds were fabricated using a custom-built near-collector electroprinting (NCE) system based on previously reported methodologies^36, 37^. Briefly, the setup consisted of a syringe pump, syringe, nozzle, nozzle holder, conductive substrate, and an XYZ translation stage with sub-micron positioning accuracy. The conductive Au/Pd-coated glass substrate was connected to a high-voltage power supply, while the metallic nozzle remained grounded. An insulating glass capillary with a tapered tip and final nozzle opening of 3 μm was mounted to the syringe.

### 2.3. Ink Preparation

Printing inks were prepared by dissolving 12.5 wt% CA in a 1:1 (v/v) acetone/DMSO solvent mixture under vortex mixing for 1 h until homogeneous. For composite formulations, 7 wt% LS was added to the CA solution and mixed under identical conditions. Higher LS concentrations were not investigated due to increased nozzle clogging during printing.

### 2.4. Scaffold Design and Fabrication

Scaffolds were designed as orthogonally patterned grid structures composed of perpendicular fibers deposited in the X and Y directions. To investigate the effect of scaffold architecture on cell behavior, grid geometries with variable pitch (center-to-center fiber spacing) ranging from 20 × 20 µm² to 250 × 250 µm², including both square and rectangular geometries, were fabricated. To systematically vary scaffold height, constructs with different numbers of printed layers were produced. Low-layer scaffolds consisting of 5 printed layers (∼5 µm total height) and high-layer scaffolds consisting of up to 80 printed layers (∼80 µm total height) were fabricated for subsequent analysis.

Scaffolds were fabricated using a custom-built NCE system following previously reported methodologies^36, 37^. Briefly, the initial nozzle-to-substrate distance was set to 10 µm, and electrohydrodynamic jetting was initiated between a grounded nozzle and a conductive substrate. Multilayer scaffolds were fabricated by sequential deposition of the designed pattern with 1 µm increments in the Z direction after each layer to achieve controlled vertical build-up. Printing was performed using glass capillaries with an inner diameter of 3 µm and an outer diameter of 9 µm. For composite scaffold fabrication, CA–LS inks were used to print 20-layer grid structures with a 10 × 20 µm pitch.

### 2.5. Scaffold Morphological Characterization

Scaffold morphology and microstructure were characterized by scanning electron microscopy (SEM; LEO 1550, Zeiss) using a secondary electron detector. Prior to imaging, samples were sputter-coated with a thin Au/Pd layer (Polaron sputter coater) to improve electrical conductivity.

### 2.6. hMSC Culture and Seeding

hMSCs were cultured in expansion medium consisting of low-glucose DMEM supplemented with 10% FBS and 1% penicillin–streptomycin. Cells were maintained at 37 °C in a humidified atmosphere containing 5% CO₂ and used at passage 4 for all experiments.

Printed scaffolds were fixed onto coated glass substrates, sterilized by UV irradiation for 15 min, and placed in 24-well plates. Prior to cell seeding, scaffolds were pre-incubated in expansion medium for at least 2 h. hMSCs were seeded onto each scaffold at a density of 30,000 cells per sample. To promote initial cell attachment, scaffolds were maintained under low-volume conditions during the first 90 min of seeding by gradually adding expansion medium in 20 μL increments every 15 min. After the adhesion period, additional medium was added to fully immerse the samples for subsequent culture.

### 2.7. LIVE/DEAD Cytocompatibility Assessment

Cell viability was assessed after 24 h of culture using a LIVE/DEAD Viability/Cytotoxicity Kit according to the manufacturer’s protocol. Briefly, culture medium was removed and samples were rinsed with sterile PBS prior to incubation in staining solution containing 0.5 μL mL⁻¹ calcein AM and 2 μL mL⁻¹ ethidium homodimer in PBS for 45 min at 37 °C in the dark. Fluorescence images were acquired using a Nikon Eclipse Ts2 fluorescence microscope. Live and dead cells were identified by green and red fluorescence, respectively. Cell viability was quantified using ImageJ as the ratio of live cells to total cells from five representative images per condition.

### 2.8. Cytoskeletal and Nuclear Staining

To visualize cell morphology and cytoskeletal organization, samples were stained for F-actin and nuclei after 24 h of culture. Cells were fixed in 4% paraformaldehyde for 30 min at room temperature, washed with PBS, and permeabilized with 0.1% Triton X-100 in PBS for 20 min. Samples were then blocked in 1% BSA for 1 h. F-actin was stained by incubating samples with Alexa Fluor 594 phalloidin (1:40 dilution in 0.1% BSA) for 45 min at room temperature. Nuclei were subsequently stained with DAPI solution (0.1% v/v in PBS) for 10 min at 37 °C. Samples were rinsed three times with PBS prior to fluorescence imaging.

### 2.9. Quantification of Nuclear Orientation

Nuclear orientation analysis was performed on low-layer scaffold conditions to quantify cellular alignment. Nuclear orientation was determined from DAPI-stained fluorescence images using ImageJ software. The angle between the major axis of each nucleus and the fiber direction was measured to assess scaffold-guided alignment. The measured angles were grouped into 10° intervals and presented as frequency distributions over a 0–180° range.

### 2.10. Quantification of Actin Coverage / Bridging Analysis

Actin coverage within high-layer scaffolds was quantified from phalloidin-stained fluorescence images using ImageJ software to assess cell bridging across scaffold openings. The actin fluorescence channel was isolated and thresholded to generate binary masks representing F-actin-positive regions. Images were segmented into predefined rectangular regions of interest (ROIs) corresponding to individual scaffold pitch units. For each ROI, the area fraction occupied by thresholded actin signal was measured and reported as actin coverage (%), representing the percentage of pore area spanned by cells. Multiple ROIs were analyzed across at least three independent fluorescence images per condition, with a minimum of three representative pore regions evaluated for each pitch size per image. Spatial heatmaps of actin coverage were generated from representative datasets to visualize local variations in cell bridging throughout multiscale scaffolds.

### 2.11. Statistical Analysis

Quantitative data are presented as mean ± standard deviation unless otherwise stated. Statistical analyses were performed using GraphPad Prism (GraphPad Software, USA). Multiple-group comparisons were analyzed using one-way analysis of variance (ANOVA) followed by Tukey’s post hoc multiple comparison test. Statistical significance was considered at *p* < 0.05.

## 3. Results and Discussion

### 3.1 Scaffold fabrication and morphology

Figure 1 shows the NCE setup together with representative scaffold architectures fabricated for this study. The printing system enabled controlled deposition of electrohydrodynamically drawn fibers into predefined multilayer grid structures (Figure 1A,B). Representative SEM images demonstrate the fabrication of scaffolds with well-defined fiber placement and regular grid architectures (Figure 1C). Alternating fiber deposition produced orthogonal scaffold geometries with uniform fiber morphology and consistent spacing. Multilayer scaffolds containing up to 80 printed layers maintained their structural integrity throughout the construct height (Figure 1D, left). High-magnification SEM images showed fiber diameters of approximately 3–5 µm (Figure 1D, right), providing the scaffold dimensions used in the subsequent cell–material interaction studies.

**Figure 1.**
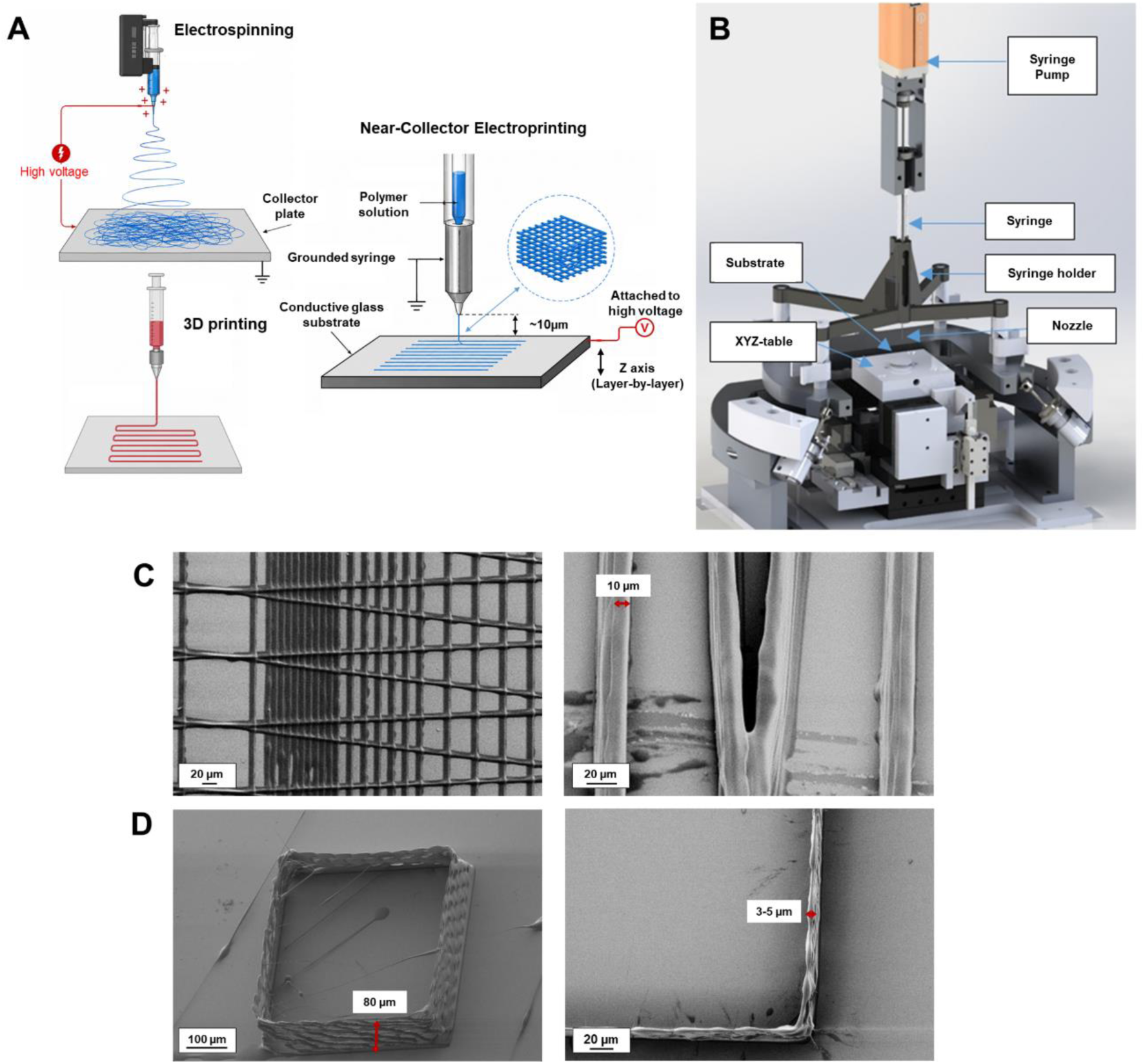
High-resolution 3D scaffold fabrication using NCE. (A) Schematic illustration of the NCE printing concept in comparison with electrospinning and extrusion-based 3D printing. NCE operates at a reduced nozzle-to-collector distance (∼10 µm), bridging the gap between random fiber formation in electrospinning and controlled fiber placement in 3D printing. (B) Schematic representation of the NCE setup used for scaffold fabrication, comprising a syringe pump, stationary syringe/nozzle assembly, conductive substrate, and computer-controlled XYZ translation stage. (C) SEM images of grid-patterned electroprinted fibers demonstrating controlled deposition of orthogonally oriented microfibers (left) and uniform fiber morphology at higher magnification (right). (D) SEM images of an 80-layer square scaffold illustrating vertical build-up capability (left) and magnified view of the scaffold edge showing stacked fibers (right). Arrows denote characteristic dimensions, including fiber diameter (∼10 µm in C, right; ∼3–5 µm in D, right) and scaffold height (∼80 µm in D, left).

### 3.2 Fabrication Fidelity, Material Versatility, and Cytocompatibility of Multiscale Ultrafine Scaffolds

#### 3.2.1 Multiscale Scaffold Design and Print Fidelity

Figure 2A illustrates the design schematic of a multiscale grid-like scaffold incorporating systematically varied pore geometries within a single construct. The design comprises orthogonally deposited fibers arranged to form systematically varied pitch sizes, ranging from 125 × 250 µm rectangular openings to 20 × 20 µm square regions.

**Figure 2.**
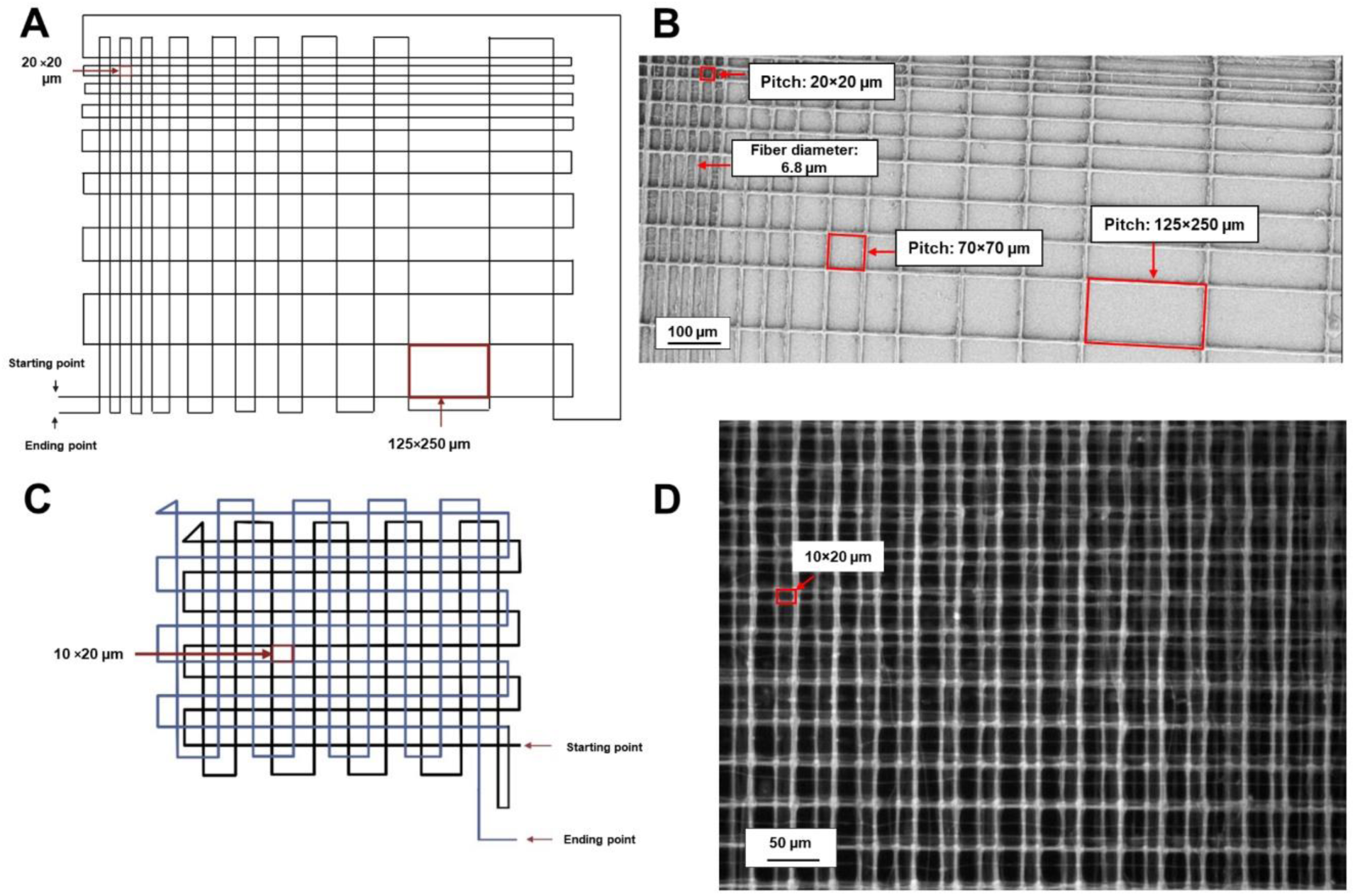
Fabrication fidelity and material versatility of multiscale ultrafine electroprinted scaffolds. (A) Schematic illustration of the multiscale scaffold design comprising orthogonally deposited fibers with systematically varied pitch sizes ranging from 125 × 250 µm to 20 × 20 µm. (B) SEM image of the fabricated multiscale CA scaffold with well-defined pitch transitions. (C) Schematic illustration of the printing strategy used to generate grid structures with feature sizes down to 10 × 20 µm, which was applied for printing CA–LS composites. (D) Optical microscopy image of a representative electroprinted CA–LS composite scaffold demonstrating retention of patterned architecture after incorporation of LS into the printing ink. Red rectangles highlight representative regions with different pitch sizes, while arrows indicate fiber diameter and spacing (pitch). Scale bars: 100 µm (B) and 50 µm (D). Panels A and C were adapted from Rezaei et al.^37^ under the Creative Commons Attribution 4.0 International (CC BY 4.0) license, with modifications.

As shown in Figure 2B, the printed construct closely reproduced the intended multiscale geometry, preserving the designed pitch transitions throughout the scaffold. High-magnification SEM imaging further confirmed successful fabrication of densely packed regions with fiber spacings down to 20 µm while maintaining structural continuity and multilayer integrity (Figure S1). Minor local deviations in fiber placement were observed in the tightest pitch regions and near turning points of the print path. These deviations could be attributed to the relatively high printing speed used (3 mm s^-1^), which may reduce jet stability during the fiber deposition. Additionally, as insulating polymer layers accumulate during the multilayer printing process, electrostatic interference may increase, potentially disrupting the electric field and affecting print fidelity. Nevertheless, the overall structural fidelity remained high across the full range of designed feature sizes.

#### 3.2.2 Material Versatility via CA–LS Composite Electroprinting

To assess whether the printing approach could accommodate composite formulations containing functional additives, LS was incorporated into the CA ink and processed under electroprinting conditions. The printing strategy is illustrated in Figure 2C, showing the toolpath and layer stacking used to generate grid structures with feature sizes of 10 × 20 µm. Optical microscopy of the resulting CA–LS constructs confirmed successful deposition of patterned multilayer architectures with preserved grid geometry (Figure 2D), demonstrating that NCE can accommodate composite formulations beyond neat polymer solutions. LS was selected as a representative functional additive due to the bioactive properties commonly associated with lignin-derived materials. Higher LS contents resulted in increased nozzle clogging during printing; therefore, a loading of 7 wt% LS was selected for subsequent experiments.

#### 3.2.3 Cytocompatibility of Printed Scaffolds

The cytocompatibility of the printed scaffolds was assessed by LIVE/DEAD staining of hMSCs cultured on CA constructs (5 layers) and CA–LS constructs (20 layers) for 24 h. Cells seeded on multiscale CA scaffolds remained predominantly viable across all tested pitch regions (35, 60, and 110 µm), with minimal red fluorescence indicating dead cells (Figure 3A–B). Quantitative analysis revealed an overall viability of 97.7 ± 1.5%, confirming that the electroprinted CA scaffolds support short-term cell survival. Similarly, CA–LS composite scaffolds maintained high cell viability (97.0 ± 0.2%), comparable to neat CA controls (Figure 3C–D), indicating that LS incorporation at the tested concentration did not adversely affect cytocompatibility. Comparable viability was also observed on multilayer square-patterned scaffolds (Figure S2), supporting the general cytocompatibility of the electroprinted structures across multiple scaffold geometries.

**Figure 3.**
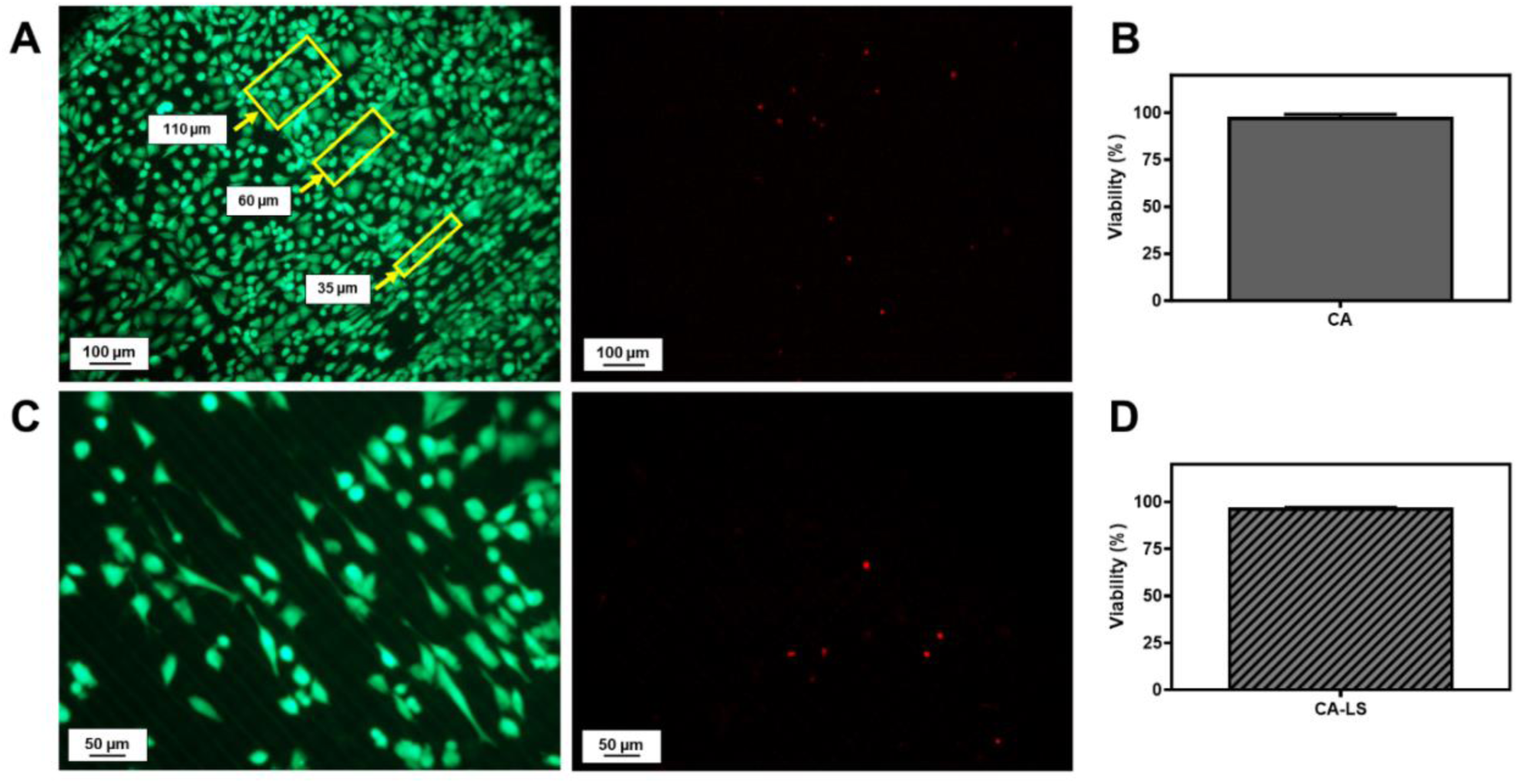
Cytocompatibility of ultrafine electroprinted scaffolds. (A) LIVE/DEAD fluorescence imaging of hMSCs cultured for 24 h on 5-layer CA scaffolds, showing high cell viability across regions with varying fiber spacing (approximately 35, 60, and 110 µm, indicated by yellow rectangles). Live cells are shown in green (left) and dead cells in red (right). (B) Quantification of hMSC viability on CA scaffolds after 24 h of culture. (C) LIVE/DEAD fluorescence imaging of hMSCs cultured for 24 h on CA–LS composite scaffolds, demonstrating high cell viability. (D) Quantification of hMSC viability on CA–LS scaffolds after 24 h of culture. Scale bars: 100 µm (A) and 50 µm (C).

### 3.3 Low-Layer Scaffolds Provide Quasi-2D Topographical Guidance

Multiscale low-layer NCE scaffolds incorporating varying fiber spacing within a single construct were used to investigate how microscale architecture influences hMSC behavior and morphology (Figure 4A). Following 24 h of culture, cytoskeletal and nuclear staining using phalloidin (actin, red) and DAPI (nuclei, blue) was performed to visualize cell alignment, spreading, and morphology in response to the defined fiber architectures (Figure 4B–D). Cells were predominantly observed on the underlying substrate rather than on the printed fibers, indicating preferential adhesion to the coated glass surface. Cellular morphology and organization were strongly dependent on local fiber spacing. In regions with relatively large spacing (∼100 µm), cells exhibited broad spreading with limited directional alignment (Figure 4B). In contrast, reducing the fiber spacing to ∼20 µm promoted pronounced elongation and alignment of both nuclei and actin filaments along the fiber direction (Figure 4C). In multiscale regions containing varying pitch values, spatially heterogeneous cell behavior was observed, with denser fiber regions supporting aligned, anisotropic morphologies and wider regions exhibiting more isotropic spreading (Figure 4D). These observations indicate that the low-layer scaffold primarily provides quasi-2D topographical cues. With a total thickness of approximately 5 µm (five layers), the structure remains small relative to the characteristic size of hMSCs, which are typically on the order of tens of micrometers^31^. Consequently, the printed fibers act as surface-guiding features rather than a volumetric 3D environment, directing cell alignment and morphology through contact guidance. These observations are consistent with previous studies demonstrating that topographic cues can regulate cell organization independently of soluble biochemical signals, with 2D substrates primarily supporting monolayer behavior^31, 40^. At the cellular level, such responses are mediated by mechanosensing pathways in which cells interpret physical cues through cytoskeletal tension and focal adhesion dynamics, leading to coordinated changes in cell polarization and nuclear organization^41^. Furthermore, engineered micro- and nanoscale architectures have been shown to control both cell and nuclear morphology through spatial confinement and directional guidance^42^. Together, these results confirm that low-layer NCE scaffolds function as quasi-2D topographical platforms, where modulation of local fiber spacing enables spatial control over cellular organization within a single construct.

**Figure 4.**
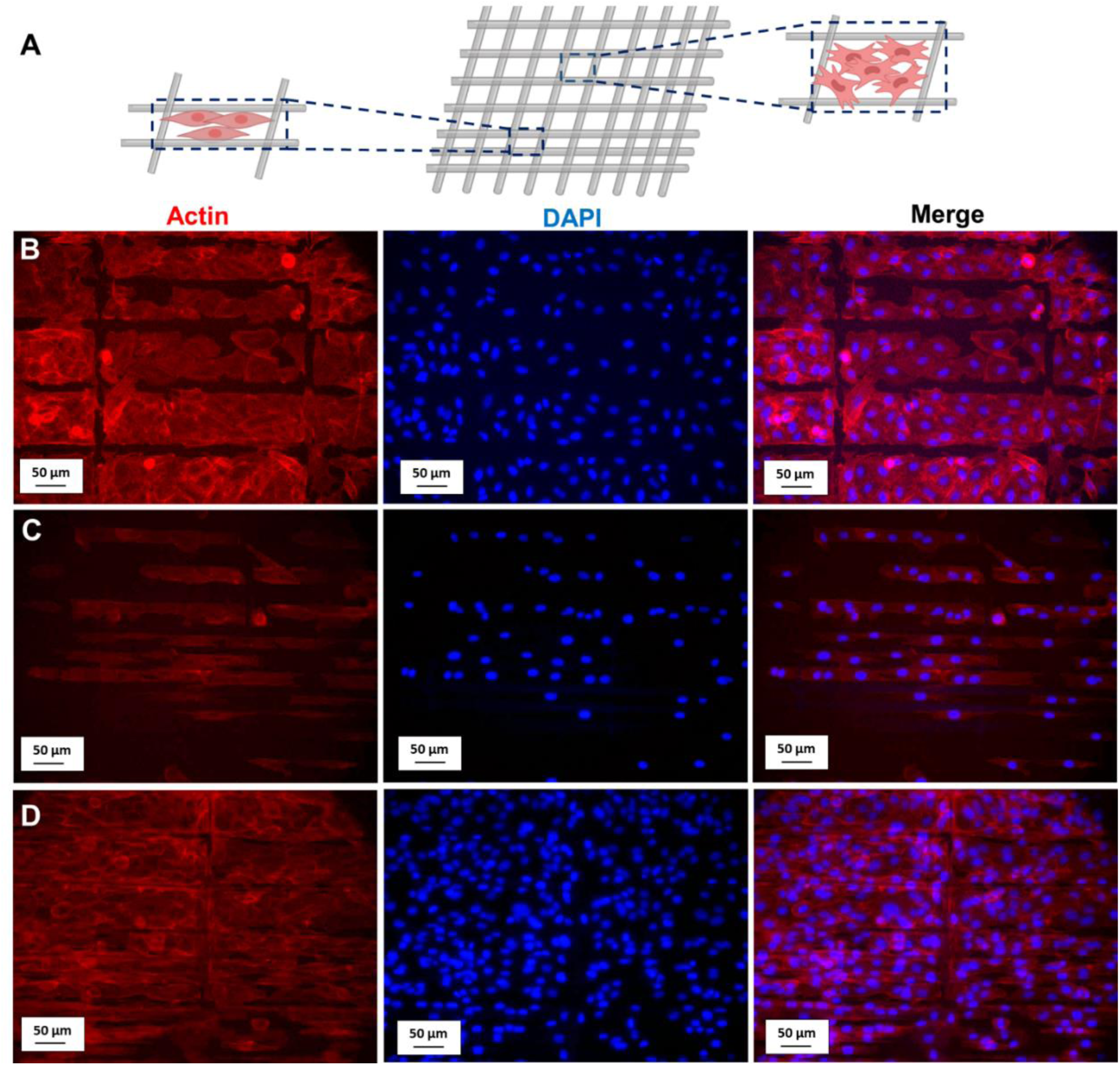
Multiscale electroprinted scaffolds guide cell morphology via quasi-2D topographical cues. (A) Schematic illustration of the experimental setup showing a low-layer (5-layer) electroprinted scaffold with varying fiber spacings. (B–D) Fluorescence microscopy images of hMSCs stained for actin filaments (red) and nuclei (DAPI, blue) after 24 h of culture, revealing distinct cellular responses to local scaffold architecture. (B) In regions with large fiber spacing (∼100 µm), cells exhibit broad spreading with limited directional organization. (C) In regions with closely spaced fibers (∼20 µm), cells align and elongate along the printed fiber direction. (D) In multiscale regions containing a gradient of fiber spacings, cells display spatially distinct morphologies, with increased alignment in densely packed regions and more isotropic spreading in wider regions. Scale bars: 50 µm.

### 3.4 Quantitative Nuclear Orientation Analysis

To quantitatively assess spacing-dependent cell alignment, nuclear orientation was analyzed relative to the printed fiber axis using ImageJ (Figure 5). Scaffolds with 100 µm fiber spacing displayed a broad and nearly isotropic orientation distribution, with no dominant alignment direction observed (Figure 5A). The corresponding DAPI image shows randomly oriented nuclei with diverse orientation angles, indicating limited topographical guidance at this spacing. In contrast, reducing the fiber spacing to 20 µm resulted in pronounced directional alignment, with 78% of nuclei oriented within 0–20° or 160–180° relative to the printed fiber direction (Figure 5B). This narrow angular distribution confirms strong unidirectional alignment along the printed scaffold architecture. This result is visually corroborated by the inset fluorescence image (DAPI staining), which highlights the elongated and uniformly aligned nuclei within the patterned region. Scatter plots of individual nuclear angles are provided in Figure S3.

**Figure 5.**
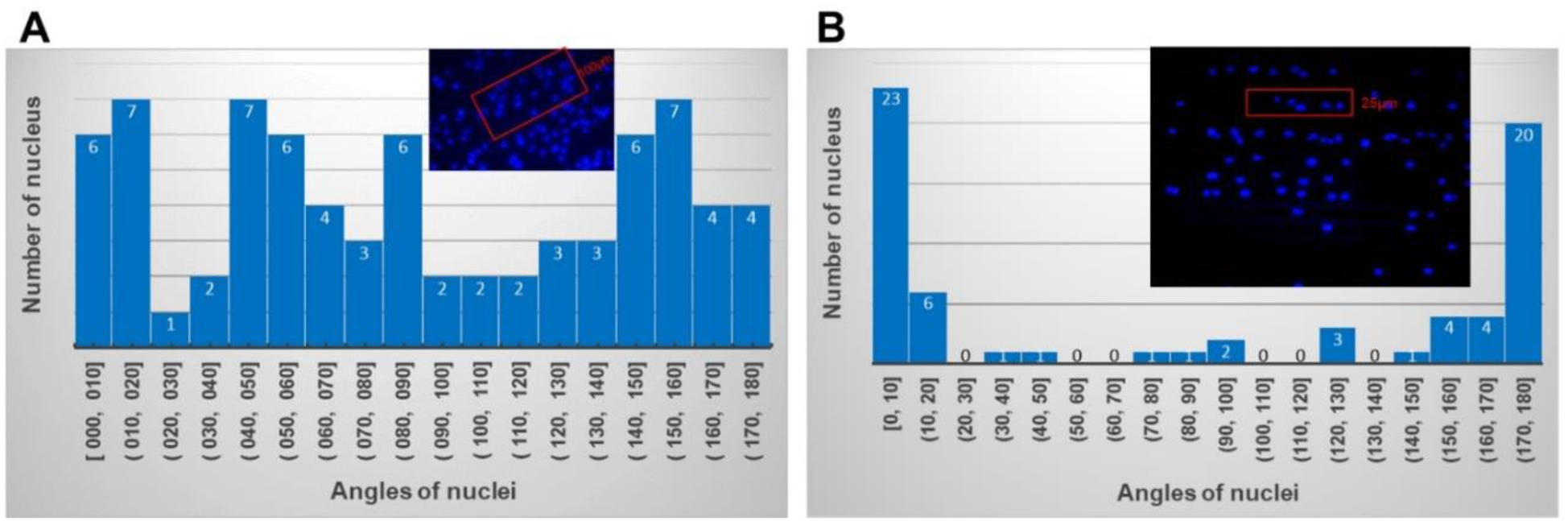
Quantitative analysis of nuclear orientation reveals spacing-dependent cellular alignment. (A) Histogram showing the distribution of nuclear orientation angles relative to the printed fiber axis for cells cultured on scaffolds with 100 µm fiber spacing. The corresponding inset image shows a broad angular distribution and reduced directional alignment, consistent with more isotropic nuclear orientation. (B) Histogram showing the nuclear orientation distribution for cells cultured on scaffolds with 20 µm fiber spacing. The inset DAPI image highlights elongated and aligned nuclei oriented along the printed fiber direction. Red rectangles indicate representative regions of interest corresponding to the local fiber spacing used for nuclear orientation analysis.

### 3.5 High-Layer Scaffolds Promote 3D Cell–Scaffold Interactions

To investigate how increased scaffold height influences cell–material interactions, hMSCs were cultured on high-layer electroprinted scaffolds consisting of 80 deposited layers (approximately 80 µm total height). A schematic illustration of the scaffold architecture and expected cell–scaffold interaction mode is shown in Figure 6A, while fluorescence images acquired after 24 h of culture are presented in Figure 6B–D. Three scaffold regions with distinct pitch values (250, 125, and 50 µm) were analyzed to assess the influence of local architecture on cellular organization. In contrast to low-layer scaffolds, cells predominantly attached to the printed fibers rather than the underlying substrate, indicating enhanced cell–scaffold interaction. Brightfield imaging shortly after seeding further confirmed that cells settled within the scaffold volume and attached between printed fibers (Figure S4).

**Figure 6.**
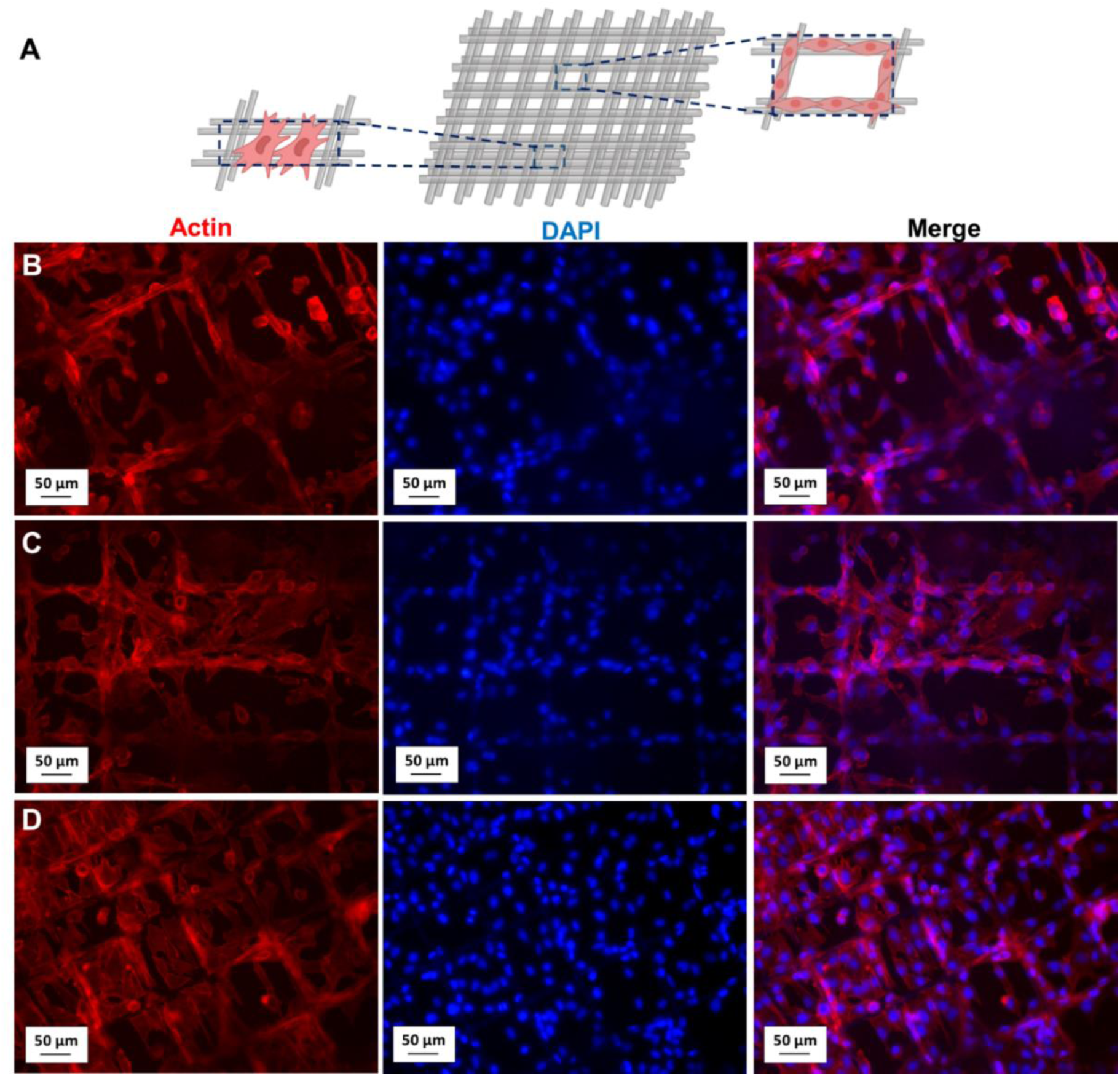
Cell morphology and cytoskeletal organization on high-layer electroprinted scaffolds with varying fiber spacings. (A) Schematic representation of an NCE scaffold consisting of 80 printed layers (∼80 µm total height), illustrating the 3D fiber grid and potential cell attachment/bridging within the scaffold architecture. (B–D) Fluorescence microscopy images of hMSCs cultured on high-layer scaffolds with varying fiber spacing. Actin filaments are stained in red and nuclei with DAPI blue; merged images are shown in the right column. (B) At 250 µm fiber spacing, cells predominantly adhered along individual fibers with limited bridging across scaffold openings. (C) At 125 µm fiber spacing, cells exhibited increased spreading and inter-fiber bridging, forming a more interconnected morphology. (D) At 50-100 µm fiber spacing, extensive cytoskeletal extensions and dense bridging between adjacent fibers were observed, indicating enhanced cellular interaction with the 3D scaffold architecture. Scale bars: 50 µm (B–D).

In regions with 250 µm spacing, cells were primarily localized along individual fibers, with limited bridging across scaffold openings (Figure 6B). Reducing the pitch to 125 µm promoted inter-fiber connectivity, with visible actin-rich extensions spanning adjacent fibers and forming a more interconnected cellular network (Figure 6C). This behavior became more pronounced at 50– 100 µm spacing, where extensive bridging and dense cytoskeletal organization were observed throughout the scaffold architecture (Figure 6D). These observations indicate that increasing structural height fundamentally alters the mode of cell–material interaction, shifting from substrate-dominated adhesion to direct engagement with the printed fiber network. Cells preferentially attach along individual fibers at large pore sizes, whereas reduced fiber spacing promotes the formation of actin-rich extensions spanning adjacent filaments, leading to interconnected cellular networks. This architecture-dependent behaviour is consistent with previous reports on microscale scaffold architectures, where structure-induced cell organization arises from fiber-guided attachment and spatial confinement effects^29^. However, the present work uniquely reveals a transition between topographical guidance and 3D bridging behavior governed jointly by scaffold height and pore geometry.

### 3.6 Quantitative Bridging Analysis

To quantitatively assess architecture-dependent bridging behavior, actin-stained fluorescence images were analyzed using a ROI-based segmentation approach, in which individual scaffold pores were evaluated for cellular coverage (Figure 7). The analyzed representative multiscale scaffold comprised a grid of systematically varied pore geometries across the construct. In the segmented images, red regions correspond to actin-positive cellular coverage within each pore (Figure 7A). The corresponding heatmap (Figure 7B) reveals a clear spatial trend in bridging behavior, with higher cellular coverage observed in regions containing smaller pore dimensions and progressively reduced coverage in larger scaffold openings.

**Figure 7.**
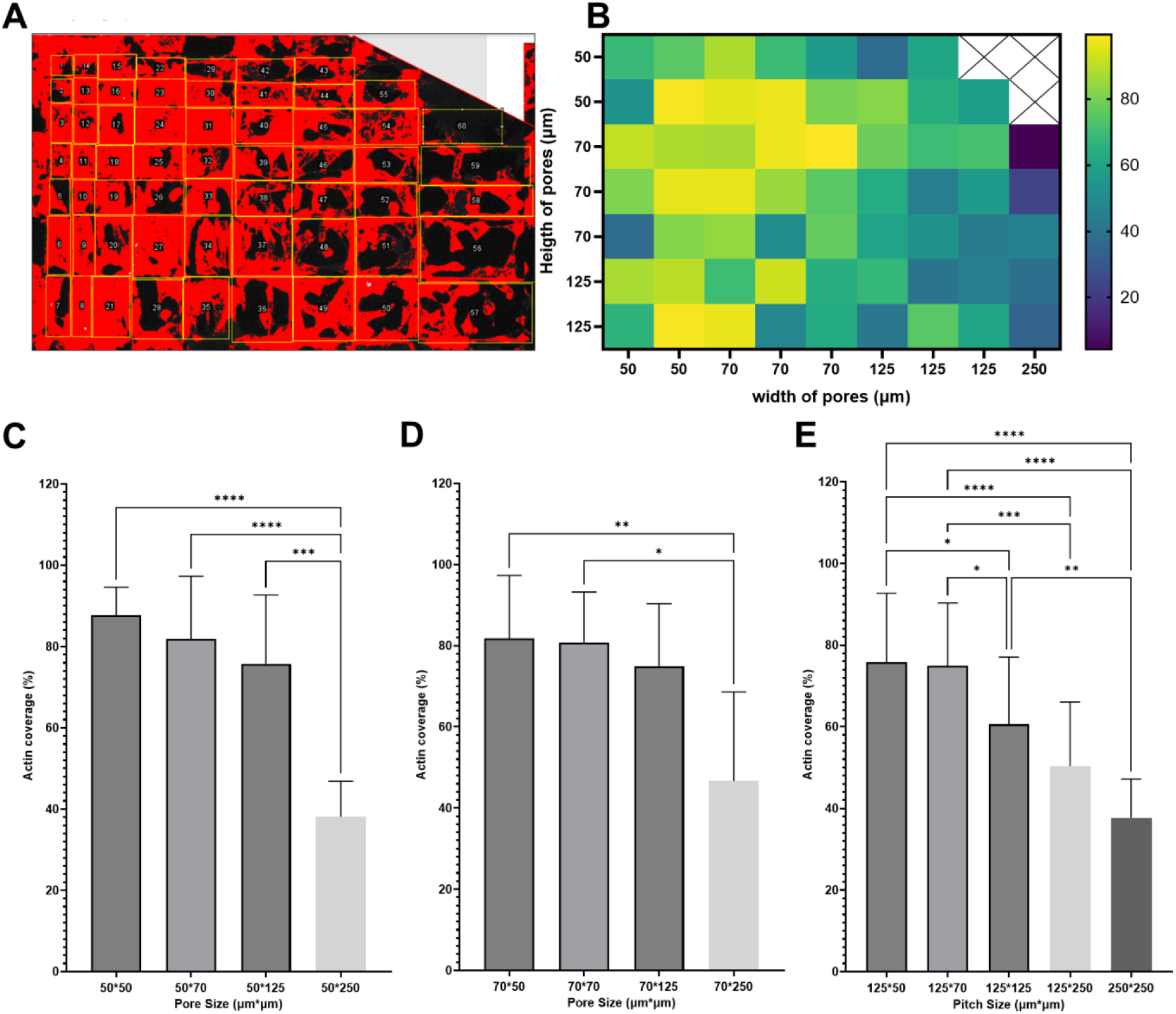
Quantitative bridging analysis reveals architecture-dependent cell–scaffold interactions in high-layer electroprinted scaffolds. (A) Representative ROI-based image analysis workflow used to quantify cellular bridging across scaffold pores from actin-stained fluorescence images. Individual pore regions were segmented and analyzed for actin-positive coverage. (B) Representative heatmap of quantified actin coverage across a multiscale scaffold, illustrating spatial variation in bridging behavior as a function of pore geometry. Cross-hatched regions indicate scaffold geometries not included in the analysis. (C–E) Quantitative analysis of bridging coverage for architectures with varying pore dimensions, grouped by fixed pore width. Bars represent mean actin coverage (%) within scaffold pores ± SD. (C) Fixed pore width of 50 µm with varying pore lengths. (D) Fixed pore width of 70 µm with varying pore lengths. (E) Fixed pore width of 125 µm with varying pore lengths, including the 250 × 250 µm condition.

Quantitative analysis confirmed that bridging coverage was strongly dependent on scaffold pore dimensions (Figure 7C–E). For scaffolds with a fixed pore width of 50 µm, bridging remained high for pore lengths of 50–125 µm (87.8 ± 6.8% and 75.8 ± 17.0%, respectively), but decreased significantly at 250 µm length to 38.1 ± 8.8% (Figure 7C). Similarly, for scaffolds with a fixed pore width of 70 µm, bridging remained high for pore lengths up to 125 µm (∼75–82%) but decreased markedly at 250 µm to approximately 47% (Figure 7D). A similar architecture-dependent decline was observed for wider scaffold openings (Figure 7E). For scaffolds with a pore width of 125 µm, bridging remained relatively high for shorter pore lengths of 50–70 µm (75.8 ± 17.0% and 74.9 ± 15.4%, respectively), but decreased progressively with increasing pore length to 60.6 ± 16.4% at 125 µm and 50.4 ± 15.7% at 250 µm. This trend culminated in a further reduction to 37.7 ± 9.5% for the largest 250 × 250 µm geometry, indicating a marked loss of bridging efficiency at larger pore dimensions. These results suggest the presence of an architectural threshold that governs the ability of hMSCs to bridge adjacent scaffold fibers within the examined culture period. When pore dimensions exceed this threshold, the distance between fibers likely surpasses the effective range of cell protrusions and cytoskeletal extension, limiting the formation of stable inter-fiber attachments. In contrast, smaller pore sizes enable cells to simultaneously engage multiple fibers, facilitating the development of tension-bearing cytoskeletal networks and promoting interconnected cellular organization. Therefore, pore geometry directly regulates the transition from isolated fiber attachment to coordinated 3D network formation. This establishes it not only as a structural parameter but also as a functional design variable, defining a quantitative window for scaffold architectures that support efficient volumetric cell integration.

## 4. Conclusion

This study demonstrates how microscale scaffold architecture governs cell–material interactions across distinct dimensional regimes using NCE cellulose scaffolds. By systematically varying scaffold layer number and pore dimensions, low-layer constructs functioned primarily as quasi-2D topographical substrates, directing cell alignment and morphology through microscale contact guidance, whereas high-layer scaffolds supported 3D cell–scaffold interactions characterized by inter-fiber bridging. Quantitative analysis further identified pore dimension as a key architectural determinant regulating bridging efficiency. In addition, incorporation of LS into the CA formulation demonstrated that composite scaffolds could be fabricated while maintaining structural integrity and cytocompatibility, highlighting the material versatility of the fabrication approach. Scaffold designs incorporating spatial variations in fiber arrangement and pore size may enable gradient tissue engineering applications requiring region-specific cellular organization. Future studies may expand this framework toward multifunctional material systems and application-specific scaffold designs for advanced tissue engineering and regenerative medicine.

## Acknowledgements

The author thanks Prof. Martin Stoddart (AO Research Institute, Davos, Switzerland) for providing the hMSCs used in this study. The author also thanks Prof. Stefan Johansson and Dr. Farnaz Rezaei for the development of the original NCE methodology on which this study was based and for permission to use representative SEM images from previous electroprinting work. The author further acknowledges Dr. Oommen Varghese for providing laboratory facilities and materials.

## Ethics statement

The hMSCs used in this study were provided by Prof. Martin Stoddart (AO Research Institute, Davos, Switzerland). Ethical approval was not required, as the cells constituted anonymized biological material and therefore fall outside the scope of the Swiss Human Research Act.

## Data availability statement

The data that support the findings of this study are available within the article and its Supplementary Information files. Additional data are available from the corresponding author upon reasonable request.

## Authorship Contributions

Mahsa Jamadi Khiabani: Conceptualization, Investigation, Formal analysis, Data curation, Visualization, Writing – original draft, Writing – review & editing.

