## Supplementary Information for "Architecture-Dependent Transition from Quasi-2D to 3D Cell–Scaffold Interactions in Ultrafine Electroprinted Cellulose Scaffolds"

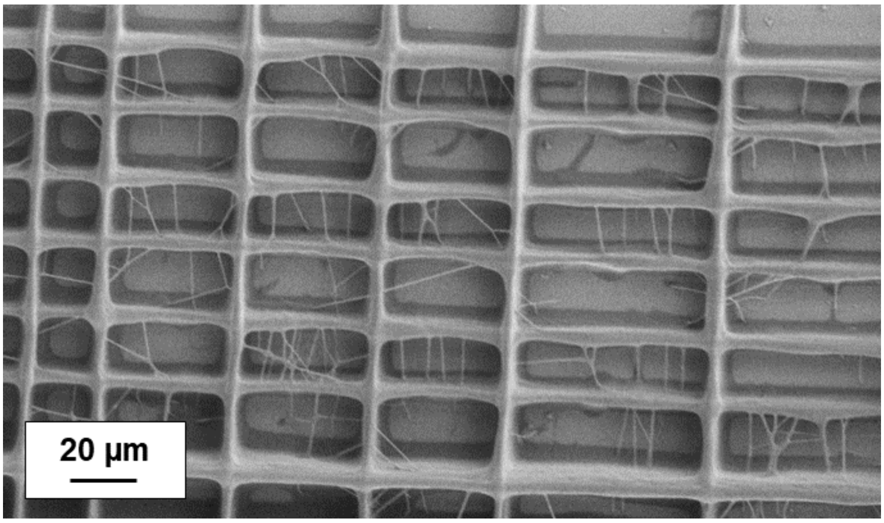


**Figure S1.** High-magnification SEM image of an ultrafine electroprinted scaffold architecture. The image shows densely packed grid structures with fiber spacing down to 20 µm, illustrating controlled fiber deposition, structural continuity, and multilayer integrity at small feature sizes. Scale bar: 20 µm.

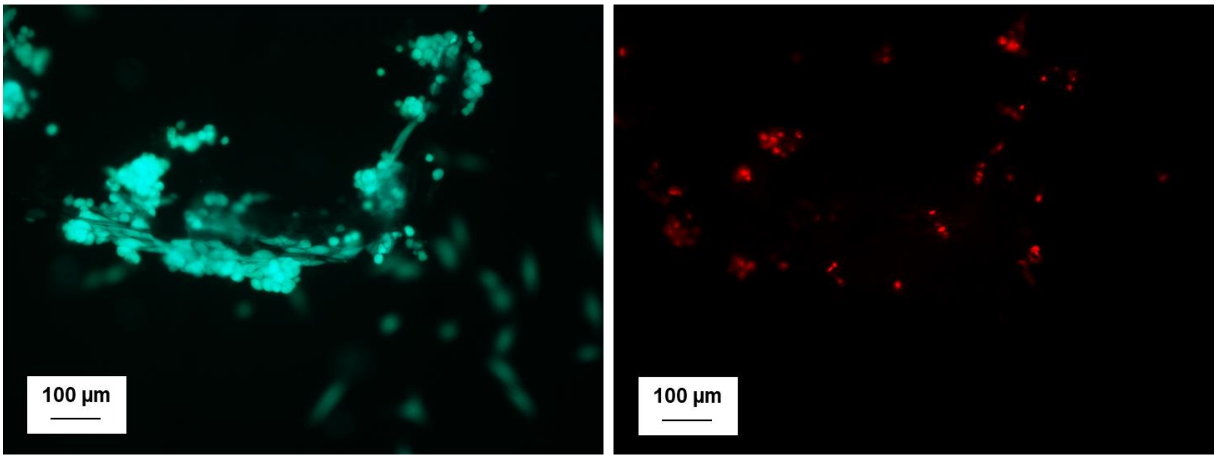


**Figure S2.** Fluorescence microscopy images of hMSCs cultured for 24h on an 80-layer NCE square scaffold. **(**Left**)** Live cells (green) indicate good scaffold biocompatibility and show alignment along the printed fibers, suggesting scaffold-guided attachment. **(**Right**)** Dead cells (red) are sparsely distributed within the scaffold. Scale bars: 100 µm.

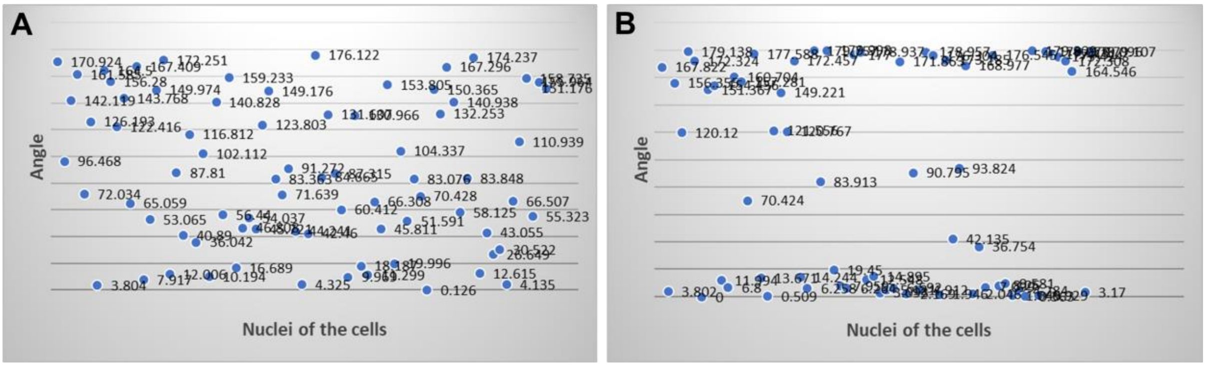


**Figure S3.** Scatter plots of individual nuclear orientation angles used for quantitative analysis. (A) Nuclear orientation in hMSCs cultured on scaffolds with a fiber pitch of 100 µm, showing a broad angular distribution. (B) Nuclear orientation in hMSCs cultured on scaffolds with a fiber pitch of 20 μm, showing preferential alignment along the fiber direction.

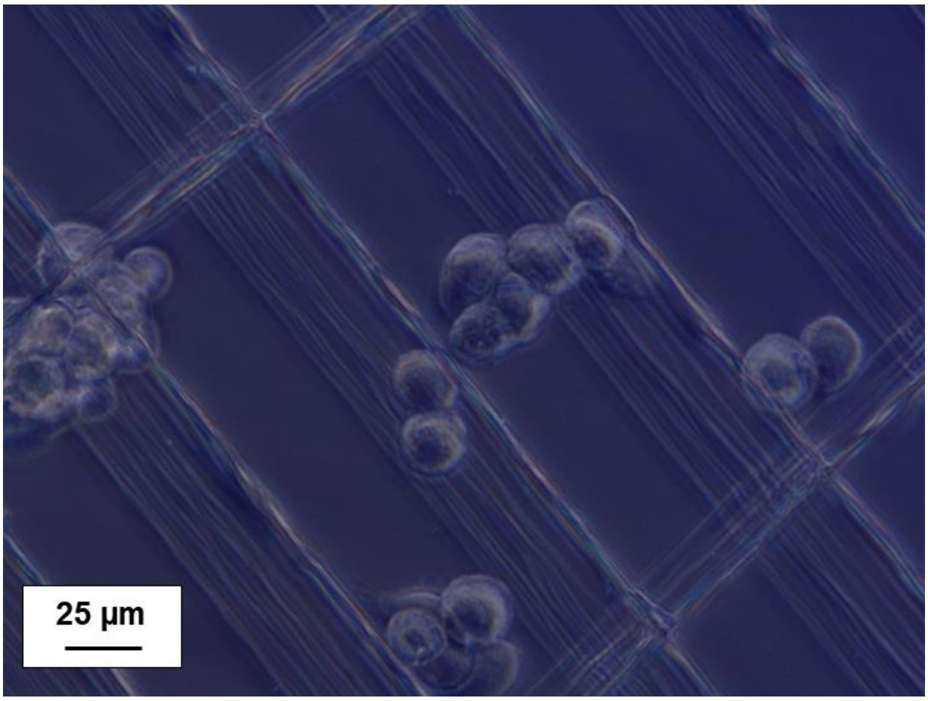


**Figure S4.** Brightfield microscopy image of hMSCs shortly after seeding onto an 80-layer NCE scaffold. Cells are visible within the scaffold architecture and between the printed fibers, illustrating their initial distribution within the structure. Scale bar: 25 μm..
